# UCSC Cell Browser: Visualize Your Single-Cell Data

**DOI:** 10.1101/2020.10.30.361162

**Authors:** Matthew L Speir, Aparna Bhaduri, Nikolay S Markov, Pablo Moreno, Tomasz J Nowakowski, Irene Papatheodorou, Alex A Pollen, Lucas Seninge, W James Kent, Maximilian Haeussler

## Abstract

**Summary:** As the use of single-cell technologies has grown, so has the need for tools to explore these large, complicated datasets. The UCSC Cell Browser is a tool that allows scientists to visualize gene expression and metadata annotation distribution throughout a single-cell dataset or multiple datasets.

**Availability and implementation:** We provide the UCSC Cell Browser as a free website where users can explore a growing collection of single-cell datasets and a freely available python package for scientists to create stable, self-contained visualizations for their own single-cell datasets. Learn more at https://cells.ucsc.edu.

## Background

Single-cell RNA-seq assays allow for the exploration of gene expression data at unprecedented detail, for surveying cellular diversity in organs (Sorragi et al. 2020) or characterizing cellular states in development (de Soysa et al. 2019) and disease (Schirmer et al. 2019). As a result, the number of publications using a single-cell RNA-seq assay has grown exponentially since 2010 (Figure S1). This growth has created a need for interactive tools that allow scientists to explore and make sense of these complex datasets at a high-level before diving deeper into computational analysis with collaborators.

Analysis of single-cell datasets typically begins with metadata and an expression matrix, which are then taken through a few standard steps: (1) normalization, (2) dimensionality reduction, (3) clustering, (4) marker gene identification and cluster labeling, and (5) visualization and sharing. Expression matrices are often normalized, trimmed, or batch-corrected. They are mapped from many dimensions to just two as x, y coordinates by dimensionality reduction algorithms like tSNE (van der Maaten. 2008) and UMAP (Dorrity et al. 2020). These algorithms attempt to retain much of the original structure. The expression matrix is also fed to a clustering algorithm in an effort to find groups of similar cells. Clusters are usually manually annotated using known cell type markers. In most research groups, cluster annotation happens by visualizing dimensionality reduction plots of well-known marker genes in the UCSC Cell Browser, Seurat, Scanpy, or one of many other tools.

## Features

The UCSC Cell Browser allows scientists to visualize the output of these single-cell analysis methods. Its primary display is a two-dimensional scatter plot, most commonly the output of tSNE or UMAP dimensionality reductions. As a single dataset may produce several x, y coordinates, the user can switch between different layouts. Scientists can pan and zoom along the plane, similar to navigating within Google Maps. Users can color cells based on provided annotations (e.g. cell type, age, detected genes) or by gene expression. There are several built-in color palettes that can be selected for coloring. The display can be split into two panes offering a side-by-side view allowing comparison using different coloring based on metadata attributes or genes (View > Split; Figure 1, upper panel). Cells can be selected, either by visually identifying and selecting groups or by combining one or more metadata-based filters (Edit > Find Cells). Selected cell identifiers can be exported for use in other analyses (Edit > Export Cells).

**Figure 1:**
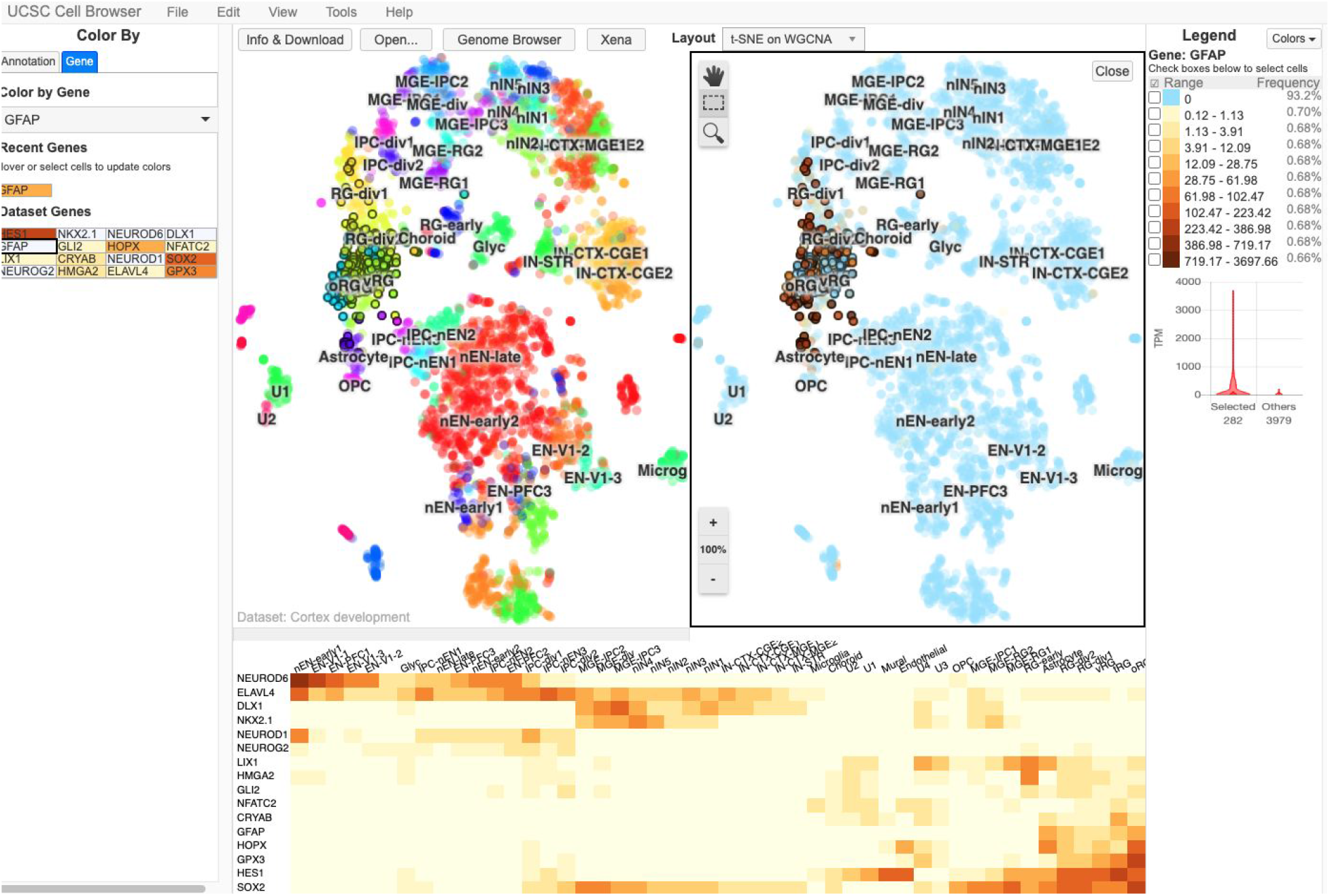
The UCSC Cell Browser interface showing a dataset focused on human cortex development. In the center of the screen, the primary layout view has been split into two vertical panes. The left pane shows the cells in this dataset colored by their metadata value for the field “WGCNACluster” (e.g. RG-div1). The right side shows the same cells colored by their expression of the gene HES1, those with a higher expression being colored dark red. On the far right side of the screen, there is a legend outlining how the colors are associated with expression bins; below this is a violin plot of this gene expression in a set of selected cells (those outlined in black in the center of the screen) versus all other cells in the dataset. On the far left side of the screen, you can see a list of metadata annotation fields available for this dataset. Each one can be clicked to color the plot by the values in that field. Below the two vertical panes, a heatmap shows 16 different genes that the authors of this dataset consider important “dataset genes”.

In addition to the primary two-dimensional scatter plot, the tool provides other ways to explore single-cell data. Datasets are typically accompanied by a list of curated “dataset genes” which are available for coloring the scatter plot. When enabled, the heatmap view shows the expression of these dataset genes across labeled clusters (View >Heatmap; Figure 1, lower panel). After selecting a group of cells, histograms showing the distribution of metadata values can be shown as well as violin plots comparing the expression of a gene in the selection against all other cells in the dataset.

The UCSC Cell Browser can further be used to display any high-dimensional data such as the bulk RNA-seq datasets from the UCSC Treehouse Childhood Cancer group (https://treehouse.cells.ucsc.edu).

## Comparison to other available tools

Data from single-cell experiments can be visualized using a number of tools in addition to the visualization options available through the analysis packages Scanpy (Wolf et al. 2018) and Seurat (Butler et al. 2018; Stuart et al. 2019) A recent bioRxiv paper (Çakır et al. 2020) compares and contrasts 13 different solutions including the UCSC Cell Browser. Features that set ours apart from these are the ability to host many datasets on a single instance arranged as a hierarchy, a simple installation procedure that requires no special server infrastructure (e.g. Flask, Shiny), and built-in converters for many data formats.

## Setting up an instance

A cell browser can be built from plain text files (expression matrix, metadata annotations, layout coordinates), but we provide utilities to import data from Seurat, Scanpy, Cellranger (Zheng, GXY et al. 2017), Loom (http://loompy.org/), and other files. Both Seurat and Scanpy provide functions to export data and build a Cell Browser instance. Installation instructions and other documentation are available at https://cellbrowser.readthedocs.io/.

The UCSC Cell Browser can be used as a Galaxy, https://galaxyproject.org/ (Afgan et al. 2018), tool for data analyzed or imported into a Galaxy instance where it has been installed. The module can also be used at the public Human Cell Atlas Galaxy instance at https://humancellatlas.usegalaxy.eu/, where it can be used to visualize reanalyzed data from the Human Cell Atlas (Regev et al. 2017) and the Single Cell Expression Atlas (Papatheodorou et al. 2020), as part of the SCAp setup (Moreno et al. 2020) which comprises more than ~90 Galaxy modules for Single Cell downstream analysis. Currently, the Bioconda (Grüning et al. 2018) module has been installed more than 1.8k times and the Galaxy module, https://toolshed.g2.bx.psu.edu/view/ebi-gxa/ucsc_cell_browser, has been cloned to ~100 Galaxy instances around the world (public and private).

## Future Plans

We have recently improved support for ATAC-seq and spatial imaging data. During the next few months, we will allow users to add custom annotations to the cells. Over the next 1-2 years, we intend to add support for running analysis algorithms (differential gene expression, dimensionality reduction, clustering) on selected cells.

## Supplemental Figures and Tables

**Figure S1:**
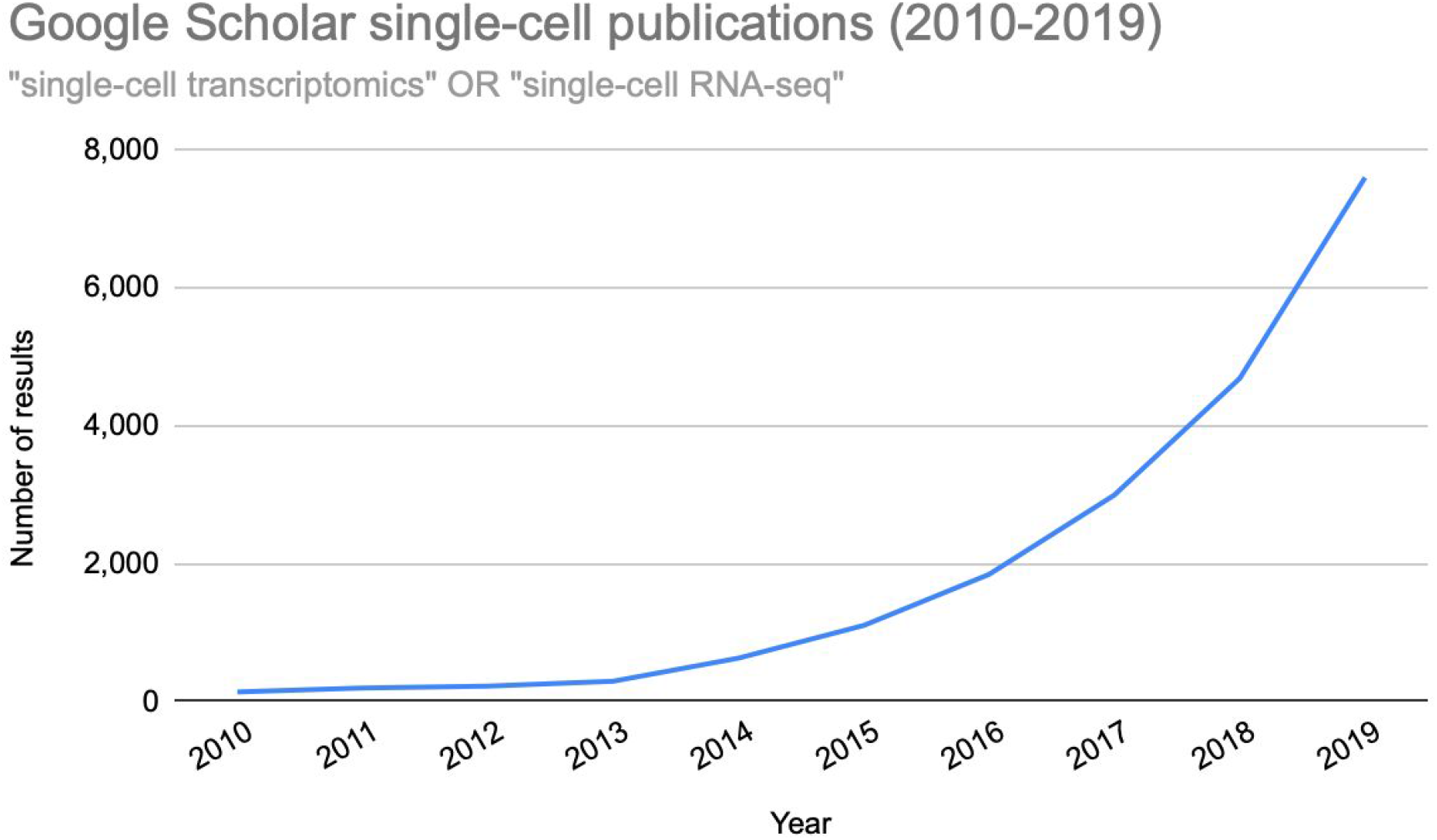
Publications matching Google Scholar search for ‘“single-cell transcriptomics” OR “single-cell RNA-seq”’ for years 2010 - 2019.

